# EPIC2: integrating single-cell chromatin accessibility with human genetics to identify trait-relevant cell types

**DOI:** 10.64898/2026.11.12.766196

**Authors:** Lingxuan Zhu, Yuchao Jiang

## Abstract

**Summary:** EPIC2 is a computational and statistical framework for prioritizing trait-relevant cellular contexts by integrating genome-wide association study (GWAS) summary statistics with single-cell chromatin accessibility data. It accounts for linkage disequilibrium when testing for association between GWAS association strength and cell-type-specific chromatin accessibility at both SNP and regulatory-peak resolutions. The resulting P-values are combined using the Cauchy combination test.

**Availability and Implementation:** The core workflow is implemented in R, with source code, vignettes, and reproducible test data freely available at https://github.com/LingxuanZ/EPIC2.

**Supplementary Information:** Supplementary material accompanies this preprint.

## 1. Introduction

Genome-wide association studies (GWAS) have identified numerous single-nucleotide polymorphisms (SNPs) significantly associated with complex traits. However, identifying the cellular contexts relevant to these associations remains challenging. Recent advances of single-cell technologies provide an unprecedented opportunity to systematically investigate the cell-type-specific enrichment of GWAS risk variants (Cuomo et al., 2023). While several methods have been developed to leverage single-cell RNA sequencing (scRNA-seq) data to prioritize trait-relevant cell types (Wang et al., 2022; Zhang et al., 2022; Yin et al., 2025), their reliance on SNP-to-gene mapping may limit their ability to capture GWAS signals from distal noncoding regulatory variants.

Single-cell assay for transposase-accessible chromatin using sequencing (scATAC-seq) measures chromatin accessibility in both coding and noncoding regions (Urrutia et al., 2019; Guan et al., 2023), providing insights into epigenetic regulation at single-cell resolution (Gaulton et al., 2023). Stratified linkage disequilibrium (LD) score regression (S-LDSC) models per-SNP heritability as a linear function of SNP-level annotations (Finucane et al., 2015). It has been used to prioritize trait-relevant cell types using annotations derived from cell-type-specific gene expression (Jagadeesh et al., 2022) or chromatin accessibility (Kim et al., 2024). S-LDSC encodes cell-type specificity through binary SNP annotations based on overlapping between SNPs and cell-type-specific gene or peak sets. These annotations, however, rely on linked gene- or peak-level measurements as proxies for SNP-level measurements and, furthermore, do not preserve quantitative differences in RNA expression or ATAC accessibility.

Here, we propose EPIC2, an LD-aware framework that prioritizes trait-relevant cell types by directly relating GWAS association strength to quantitative cell-type-specific chromatin accessibility without requiring SNP-to-gene mapping. Its central biological hypothesis is that, if a particular cell type is relevant to a trait, polygenic GWAS signals are enriched at loci preferentially accessible in that cell type. EPIC2 performs four tests at SNP and regulatory-peak resolutions, using cell-type-specific count-based accessibility and Wilcoxon rank-sum (WRS) test *Z*-scores as input. For each combination of a trait and a cell type, the resulting *P*-values are combined using a heavy-tailed combination method to obtain an omnibus *P*-value (Gui et al., 2025), allowing for downstream functional investigation in a cell-type-specific manner.

## 2. Approach and implementation

In the sections below, we outline each of four model configurations of EPIC2, while leaving the details on data preprocessing, feature construction, model fitting, and implementation in the Supplementary Methods.

### 2.1. SNP-level prioritization model

Let *j* index the SNPs for a given GWAS trait, and *k* index the focal cell-type groups. At SNP resolution, EPIC2 fits the following generalized least squares regression:

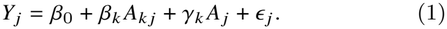

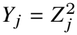 represents non-directional GWAS association magnitude at SNP *j*, where *Z* _*j*_ denotes the corresponding GWAS *Z*-score. Squaring the GWAS *Z*-score allows variants with either positive or negative allelic effects to contribute according to the strength of their association, because chromatin accessibility does not indicate the direction of these effects. Under the null hypothesis, *Y*_*j*_ follows a 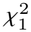 distribution. Under a bivariate normal null model, 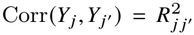 _′_, where 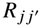 denotes the genotype correlation between SNPs *j* and *j* ^′^. The LD matrix 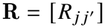 is estimated from the 1000 Genomes Project reference panel (1000 Genomes Project Consortium, 2015). Such LD-induced correlation is modeled through the residual vector ***ε***, which has mean zero and working covariance *σ*^2^***Σ***, where 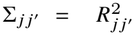 . When constructing ***Σ***, entries for SNP pairs on different chromosomes or more than 5 Mb apart are set to zero.

*A*_*kj*_ measures cell-type specificity from scATAC-seq data and takes two alternative forms: a normalized cell-type-specific ATAC measure or a cell-type-specific WRS test score. In the former case, *A*_*k j*_ is the mean of library-size-adjusted absolute accessibility counts at SNP *j* across cells of type *k*. To construct *A*_*k j*_, ATAC fragments overlapping the retained SNPs are first counted to generate a sparse SNP-by-cell count matrix, which is then corrected for the library-size factor for each cell, followed by averaging across all cells of type *k*. The baseline covariate *A* _*j*_ is the mean library-size-adjusted accessibility across all retained cells, included to adjust for overall accessibility at SNP *j* . In the WRS score specification, *A*_*k j*_ is replaced with 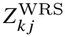, the winsorized *Z*-score from a one-sided and one-versus-rest WRS test of higher accessibility in cells of type *k* than in all other cells. The baseline covariate *A* _*j*_ remains unchanged from the count-based specification.

For both SNP-level specifications, EPIC2 tests the prespecified enrichment hypothesis for each cell-type group *k*, with *H*_0_ : *β*_*k*_ ≤ 0 against *H*_1_ : *β*_*k*_ > 0. The LD-induced working covariance matrix, ***Σ***, is transformed with a ridge-stabilized inverse square root, applied to the response, intercept, group-specific predictor, and baseline accessibility predictor for whitening. Standard errors and one-sided *P*-values are obtained from the transformed ordinary least-squares fit using its residual degrees of freedom. Details of the regularization and whitening procedure are provided in the Supplementary Methods.

### 2.2. Peak-level prioritization model

At peak resolution, let *p* index the ATAC peaks. For each cell-type group *k*, EPIC2 fits the following regression model across peaks:

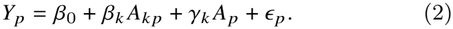

GWAS *Z*-scores are combined as 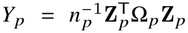, where *n* _*p*_ denotes the number of GWAS variants overlapping peak *p*, **Z**_*p*_ is the vector of GWAS *Z*-scores, and Ω_*p*_ is the ridge-regularized inverse of their within-peak LD matrix. Details of the regularization procedure are provided in the Supplementary Methods.

As in the SNP-level analysis, *A*_*k p*_ takes one of two forms: the normalized cell-type-specific accessibility of peak *p* in cell type *k*, or its corresponding cell-type-specific WRS *Z*-score. Both peak-level specifications incorporate baseline accessibility *A*_*p*_, defined analogously to *A* _*j*_, to adjust for overall accessibility at peak *p*. The term *ϵ* _*p*_ denotes residual variation in *Y*_*p*_ not explained by the focal accessibility predictor and the baseline covariate.

The working covariance between peak pairs is calculated from cross-peak LD in the reference panel for peaks on the same chromosome and within 5 Mb of each other, and is set to zero otherwise. See Supplementary Methods for details. The resulting sparse covariance matrix is regularized, and its inverse square root is used to whiten the peak-level response and full design matrix as in the SNP-level analysis. The same one-sided enrichment hypothesis is tested for *β*_*k*_ .

### 2.3. Cauchy combination test

For each pair of trait and cell type, the nominal one-sided *P*-values from the SNP-count, SNP-WRS, peak-count, and peak-WRS models are combined with equal weights using the Cauchy combination test, a heavy-tailed method for aggregating dependent *P*-values (Liu and Xie, 2020; Gui et al., 2025). The combined *P*-values are adjusted using the Benjamini-Hochberg (BH) procedure.

## 3. Prioritizing trait-relevant cell types for lipid traits and psychiatric disorders

We applied EPIC2 to a human cell atlas of fetal chromatin accessibility (Domcke et al., 2020) to prioritize trait-relevant cell types for four lipid traits and four psychiatric datasets: high-density lipoprotein cholesterol (HDL), low-density lipoprotein cholesterol (LDL), total cholesterol (TC), triglycerides (TG) (Willer et al., 2013), schizophrenia (SCZ from Schizophrenia Working Group of the Psychiatric Genomics Consortium (2014) and SCZ2 from Pardiñas et al. (2018)), bipolar disorder (BIP) (Stahl et al., 2019), and schizophrenia or bipolar disorder (SCZBIP) (Ruderfer et al., 2018).

For scATAC-seq quality control (QC), cells were filtered based on transcription start site (TSS) enrichment score, fraction of fragments within peaks, total ATAC fragment counts within peaks, nucleosome signal, and fraction of fragments overlapping blacklist regions. Cell-type groups with fewer than 50 cells were excluded, and we focused on major cell types across seven tissues: cerebrum, liver, kidney, adrenal, intestine, lung, and heart. Only autosomal peaks with lengths of 100–1,000 bp and non-zero coverage were retained for analysis. For GWAS QC, only SNPs present in the 1000 Genomes Project reference panel (1000 Genomes Project Consortium, 2015) were retained. Duplicate rsIDs and strand-ambiguous SNPs were removed. Variants were required to have a minor allele frequency of at least 0.05 in both the GWAS data and the reference panel. LD pruning was performed using SNPRelate with a threshold of 0.8. The post-QC scATAC-seq dataset contained 84,238 cells and 915,345 peaks. Trait-specific GWAS processing and final SNP counts are summarized in Supplementary Table S1. Following standard scATAC-seq analysis pipelines (Guan et al., 2023), dimension reduction of cells with tissue and cell-type annotations is visualized in Supplementary Fig. S1. As a sanity check, the top cell-type-specific SNPs and peaks from the WRS test also showed predominantly cell-type-specific accessibility patterns (Supplementary Figs. S2 and S3).

EPIC2’s Cauchy combination test identified 12 significant associations between traits and cell types across six GWAS traits after within-trait BH correction (Fig. 1). Specifically, liver hepatoblasts were prioritized for LDL, TC, and TG, while liver myeloid cells were prioritized for LDL and TC. The hepatoblast findings are consistent with both the liver’s central role in lipid metabolism at the tissue level and previously reported enrichment of lipid GWAS signals in hepatocytes at the cell-type level (Wang et al., 2022; Hao et al., 2020). In addition, kidney ureteric bud cells were prioritized for LDL and TC, and kidney vascular endothelial cells were prioritized for TG, in line with previous reports linking renal processes to lipid dysregulation (Molina-Jijon et al., 2020; Clement et al., 2014).

**Figure 1.**
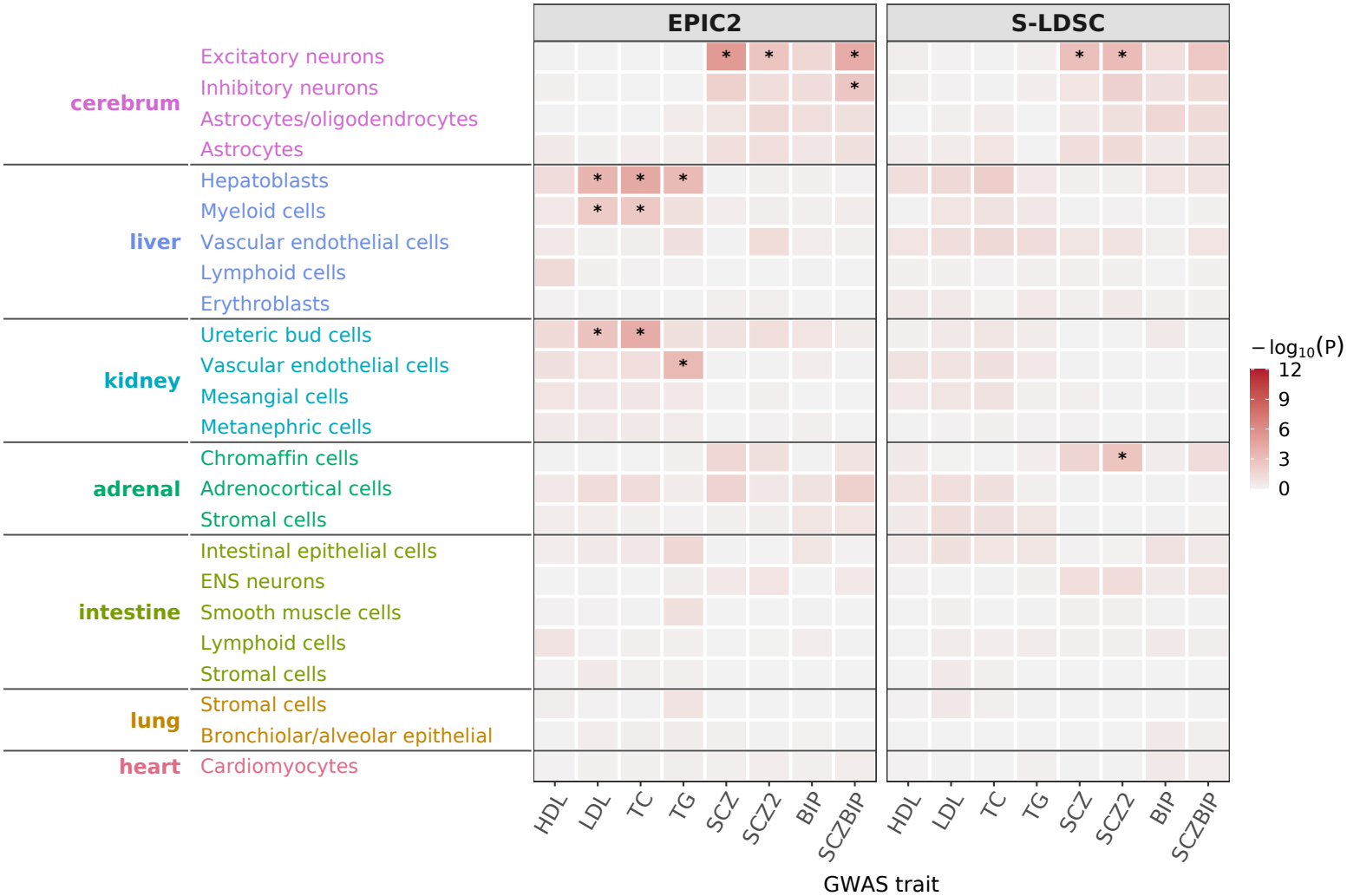
Comparison of EPIC2 and S-LDSC for cell-type prioritization across lipid traits and psychiatric disorders. Rows represent cell-type groups, and columns represent GWAS datasets. EPIC2 combines four one-sided *P*-values using the Cauchy combination test, whereas S-LDSC performs a one-sided test using cell-type-specific ATAC annotations. Tile color corresponds to − log_10_ (*P*) for unadjusted *P*-values, capped at 12. Asterisks indicate significance at a BH-adjusted *P*_adj_ < 0.05 within each GWAS trait.

For psychiatric disorders, the strongest enrichment was observed for SCZ in cerebrum excitatory neurons (*P*_adj_ = 1.30 × 10^−4^), which were also prioritized for SCZ2 and SCZBIP. Such findings are consistent with enrichment of common SCZ-associated variants in open chromatin regions of glutamatergic neurons (Hauberg et al., 2020). EPIC2 also prioritized cerebrum inhibitory neurons for SCZBIP (*P*_adj_ = 2.98 × 10^−2^). Previous work reported enrichment of SCZ genetic associations in genes preferentially expressed in specific prenatal excitatory and inhibitory neuronal populations (Cameron et al., 2023). Heritability enrichment has also been reported near genes specifically expressed in excitatory neurons for SCZ and in GABAergic neurons for BIP (Finucane et al., 2018). Detailed results from the four component models across the aforementioned 12 enrichment by EPIC2 are provided in Supplementary Table S2. Notably, TC in liver hepatoblasts was the only pair, for which all four component tests were significant after within-trait BH correction. For other combinations of traits and cell types, the four testing schemes provide complementary evidence for cell-type prioritization.

For benchmarking, we compared EPIC2 against S-LDSC (Finucane et al., 2015). Details on the construction of the cell-type-specific annotations from scATAC-seq and the S-LDSC analysis pipeline are provided in the Supplementary Methods. For cell-type prioritization, S-LDSC identified three significant associations between traits and cell types after BH correction. Both EPIC2 and S-LDSC prioritized excitatory neurons from the cerebrum for SCZ and SCZ2, while adrenal chromaffin cells were prioritized for SCZ2 only by S-LDSC. The S-LDSC result for SCZBIP in cerebrum excitatory neurons was nominally significant (*P* = 2.72 × 10^−3^), but it did not remain significant after BH correction (*P*_adj_ = 6.54 × 10^−2^). Additionally, all associations between lipid traits and cell types identified by EPIC2 had positive S-LDSC annotation coefficients, although none were statistically significant after BH correction (Supplementary Table S3). Together, EPIC2 identified associations between traits and cell types consistent with prior genetic evidence, demonstrating its outperformance for cell-type prioritization.

## 4. Conclusion

EPIC2 integrates GWAS summary statistics with cell-type-specific chromatin accessibility measures from scATAC-seq. It accounts for local LD structure across both SNPs and peaks, and performs four component tests with different resolutions and configurations, followed by an omnibus test. Through empirical analyses of a human cell atlas and eight GWAS datasets, we demonstrate that EPIC2 recapitulates known disease etiology and outperforms S-LDSC in prioritizing trait-relevant cell types.

## Acknowledgements

The authors thank High Performance Research Computing at Texas A&M University for providing computational resources and support.

## Author contributions

Lingxuan Zhu (Data Curation, Formal Analysis, Methodology, Software, Visualization, and Writing Original Draft) and Yuchao Jiang (Conceptualization, Methodology, Project Administration, Supervision, and Writing Review and Editing).

## Funding

This work was supported by the National Institutes of Health (NIH) Grant R35 GM138342 to YJ.

## Conflict of interest

None declared.

## Data availability

The fetal scATAC-seq data are available from GEO under accession GSE149683. GWAS summary statistics are available from the repositories cited in the corresponding publications. Source code, documentation, and vignettes containing non-restricted test data are available at https://github.com/LingxuanZ/EPIC2.

## Notes

https://github.com/LingxuanZ/EPIC2

https://lingxuanz.github.io/EPIC2/EPIC2_vignette.html

### Competing Interest Statement

The authors have declared no competing interest.

