## Supplementary material for "EPIC2: integrating single-cell chromatin accessibility with human genetics to identify trait-relevant cell types"

**S3 Supplementary Discussion** **8**

**S4 Supplementary Figures** **11**

**S5 Supplementary Tables** **14**

### S1 Supplementary Methods

#### S1.1 Study design and workflow

EPIC2 tests for positive associations between GWAS association strength and cell-type-specific chromatin accessibility at SNP and ATAC-seq peak resolutions, adjusting for overall accessibility and accounting for LD. The workflow includes preprocessing of fetal scATAC-seq and GWAS data, construction of trait-specific SNP- and peak-level accessibility features, and LD-aware association testing using count-based accessibility and Wilcoxon rank-sum (WRS) scores. For each combination of a trait and a cell type, EPIC2 reports four nominal one-sided component $P$-values and an omnibus $P$-value obtained by combining them with equal weights using the Cauchy combination test.

#### S1.2 Data sources

We used data from the human fetal chromatin accessibility atlas of Domcke et al. (2020) (GEO accession GSE149683), which contains high-quality chromatin accessibility profiles from 53 fetal samples representing 15 organs. The preprocessed subset analyzed in this study contained cells from seven tissues: heart, intestine, adrenal gland, liver, kidney, lung, and cerebrum. All genomic coordinates used in the analysis were referenced to the hg19/GRCh37 assembly.

Summary statistics for high-density lipoprotein cholesterol (HDL), low-density lipoprotein cholesterol (LDL), total cholesterol (TC), and triglycerides (TG) were obtained from the Global Lipids Genetics Consortium meta-analysis (Willer et al., 2013). This meta-analysis combined association results from GWAS and Metabochip studies involving up to 188,577 European-ancestry individuals. Sample sizes in the released summary statistics vary across variants.

The four psychiatric GWAS datasets were labeled SCZ, SCZ2, BIP, and SCZBIP. SCZ denotes the 2014 schizophrenia GWAS reported by the Psychiatric Genomics Consortium (PGC). The study's primary GWAS dataset comprised 49 non-overlapping case–control cohorts (46 of European and three of East Asian ancestry; 34,241 cases and 45,604 controls) and three European-ancestry family-based samples comprising 1,235 trios comprising an affected offspring and both parents (Schizophrenia Working Group of the Psychiatric Genomics Consortium, 2014). Thus, the reported sample size of 79,845 includes only case–control participants and excludes the family-based trios. SCZ2 denotes the CLOZUK–PGC2 meta-analysis of 40,675 schizophrenia cases and 64,643 controls of European ancestry (105,318 participants; Pardiñas et al. 2018). BIP denotes the discovery GWAS meta-analysis of 20,352 bipolar disorder cases and 31,358 controls of European ancestry (51,710 participants; Stahl et al. 2019). SCZBIP denotes the combined schizophrenia or bipolar disorder case–control GWAS, comprising 53,555 cases (33,426 schizophrenia and 20,129 bipolar disorder) and 54,065 controls (107,620 participants; Ruderfer et al. 2018). The two disorders were analyzed as a single case group. SCZ and SCZ2 shared PGC schizophrenia samples, while SCZBIP included samples from the PGC schizophrenia and bipolar disorder collections contributing to SCZ and BIP, respectively. These datasets are therefore not independent, and concordant findings across them should not be interpreted as independent replication.

#### S1.3 scATAC-seq preprocessin g

The preprocessed, merged Seurat object contained 87,526 cells and a shared set of 1,050,819 ATAC-seq peaks. Quality-control metrics were calculated with Signac (Stuart et al., 2021). Cells were first retained if the transcription start site (TSS) enrichment score was at least 2.5 and the fraction of fragments overlapping peaks (FRiP) was at least 0.3. The first percentile of total ATAC-seq counts within peaks was then calculated across all remaining cells, and cells with counts below this threshold were removed. Cells were further required to have a fraction of fragments overlapping blacklist regions below 0.02 and a nucleosome signal of at most 4. Cell-type groups with fewer than 50 cells after cell-level filtering were excluded.

Peaks with zero total counts across the retained cells were removed, as were non-autosomal peaks, peaks shorter than 100 bp or longer than 1,000 bp, and peaks overlapping ENCODE blacklist regions (Amemiya et al., 2019). The resulting peak-by-cell matrix contained 915,345 peaks and 84,238 cells. Term frequency–inverse document frequency (TF–IDF) normalization and singular value decomposition were performed with Signac to obtain a latent semantic indexing (LSI) representation. For quality assessment, LSI dimensions 2–30 were used for visualization with uniform manifold approximation and projection (UMAP) and shared nearest neighbor (SNN) graph-based clustering in Seurat.

#### S1.4 GWAS processing and trait-specific accessibility matrices

Variants in the aligned GWAS summary statistics were matched by rsID to variants in the European-ancestry subset of the 1000 Genomes Project Phase 3 reference panel (1000 Genomes Project Consortium, 2015). Duplicate rsID entries were removed, retaining the first occurrence. Variants were required to have a minor allele frequency (MAF) of at least 0.05 in both the GWAS data and the reference panel. Autosomal SNPs were LD-pruned chromosome by chromosome using SNPRelate with the composite LD measure, a 500-kb sliding window, an absolute LD threshold of 0.8, and a random seed of 1000. Strand-ambiguous SNPs (A/T and C/G) were subsequently removed. Unpruned aligned GWAS summary statistics were processed separately for S-LDSC. GWAS $Z$-scores used in the primary models were calculated by dividing each reported effect estimate by its corresponding standard error.

For each trait, GWAS SNPs retained after QC and LD pruning were further restricted to those located within the filtered ATAC peaks. ATAC fragments overlapping these SNP positions were counted to generate a sparse SNP-by-cell count matrix, and SNPs with zero total fragment coverage across all retained cells were removed. Separately, the existing peak-by-cell ATAC count matrix was restricted to peaks overlapping at least one GWAS SNP retained after QC and LD pruning. Thus, the two matrices quantify accessibility at retained GWAS SNP positions and across GWAS-overlapping ATAC peaks, respectively.

#### S1.5 Cell-type accessibility scores

For each trait, one-versus-rest Wilcoxon rank-sum (WRS) tests were performed for each feature in the sparse SNP-by-cell and GWAS-overlapping peak-by-cell count matrices using `presto::wilcoxauc` (Korsunsky et al., 2022). Each retained cell-type group was compared with all remaining cells in the corresponding matrix.

At SNP resolution, let $p_{2,kj}$ and $\mathrm{AUC}_{kj}$ denote the two-sided nominal $P$-value and area under the receiver operating characteristic curve for SNP $j$ and cell-type group $k$, respectively. Values of $\mathrm{AUC}_{kj} > 0.5$ indicate that accessibility values tend to be higher in cell-type group $k$ than in the remaining cells. The one-sided $P$-value for greater accessibility in cell-type group $k$ was calculated as

$$p_{kj}^{(+)} = \begin{cases} p_{2,kj}/2, & \mathrm{AUC}_{kj} \geq 0.5, \\ 1 - p_{2,kj}/2, & \mathrm{AUC}_{kj} < 0.5. \end{cases} \tag{S1}$$

The corresponding score was $Z_{kj}^{\mathrm{WRS}} = \Phi^{-1}\{1 - p_{kj}^{(+)}\}$, where $\Phi$ is the standard normal cumulative distribution function. At peak resolution, the same procedure was applied with $j$ replaced by the peak index $p$.

Within each trait and genomic resolution, these scores were winsorized at the empirical 2nd and 98th percentiles across all features and cell-type groups, with percentile cutoffs estimated from finite scores. In the association models below, $Z_{kj}^{\mathrm{WRS}}$ and $Z_{kp}^{\mathrm{WRS}}$ denote the winsorized SNP- and peak-level scores, respectively.

#### S1.6 SNP-level association models

For a given trait, GWAS association strength at SNP $j$ was represented by $Y_j = Z_j^2$, where $Z_j$ denotes the corresponding GWAS $Z$-score. Let $C_{jc}$ denote the number of ATAC fragments overlapping SNP position $j$ in cell $c$, $L_c$ the total ATAC count for cell $c$ across all retained autosomal peaks, and $s_c = L_c / \operatorname{median}_{c'} L_{c'}$ the library-size factor for cell $c$. For cell type $k$, the mean library-size-adjusted accessibility and the mean baseline accessibility across all retained cells were defined as

$$A_{kj} = \frac{1}{N_k} \sum_{c \in \mathcal{C}_k} \frac{C_{jc}}{s_c}, \qquad A_j = \frac{1}{N} \sum_{c=1}^{N} \frac{C_{jc}}{s_c}, \tag{S2}$$

where $\mathcal{C}_k$ denotes the set of retained cells belonging to cell type $k$, $N_k = |\mathcal{C}_k|$ is its cell count, and $N$ is the total number of retained cells.

For each cell type, we fitted a count-based accessibility model and a WRS-score model:

$$Y_j = \beta_0 + \beta_k A_{kj} + \gamma_k A_j + \epsilon_j, \tag{S3}$$

$$Y_j = \beta_0 + \beta_k Z_{kj}^{\mathrm{WRS}} + \gamma_k A_j + \epsilon_j. \tag{S4}$$

Here, $Z_{kj}^{\mathrm{WRS}}$ is the winsorized cell-type-specific WRS score defined in Section S1.5. The count-based baseline covariate $A_j$ was retained in both models, and coefficients were estimated separately for each specification. The term $\epsilon_j$ denotes residual variation in $Y_j$ not explained by the focal accessibility predictor and the baseline covariate. The prespecified enrichment hypothesis for both models was $H_0 : \beta_k \leq 0$ against $H_1 : \beta_k > 0$.

Pairwise Pearson genotype correlations $R_{jj'}$ were calculated from the 1000 Genomes Project reference panel (1000 Genomes Project Consortium, 2015) for retained SNP pairs on the same chromosome and within 5 Mb. Throughout, reference-panel LD estimates and the matrices derived from them were treated as fixed inputs to the working models. Under a bivariate standard normal null model for the GWAS $Z$-scores with $\operatorname{Corr}(Z_j, Z_{j'}) = R_{jj'}$, $\operatorname{Cov}(Z_j^2, Z_{j'}^2) = 2R_{jj'}^2$ and $\operatorname{Var}(Z_j^2) = 2$, giving $\operatorname{Corr}(Y_j, Y_{j'}) = R_{jj'}^2$. We therefore constructed a sparse local working matrix $\mathbf{\Sigma}$ with $\Sigma_{jj'} = R_{jj'}^2$ for SNP pairs on the same chromosome and within 5 Mb, zero otherwise, and $\Sigma_{jj} = 1$. The residual vector $\boldsymbol{\epsilon}$ was modeled as having mean zero and a working covariance given by $\sigma^2$ times a regularized version of $\mathbf{\Sigma}$, where $\sigma^2$ is the residual variance scale.

Let $b$ index the connected components defined by the nonzero off-diagonal entries of $\mathbf{\Sigma}$, with isolated SNPs treated as singleton components. The submatrix for component $b$ had eigendecomposition $\mathbf{\Sigma}_b = U_b \operatorname{diag}(d_b) U_b^{\mathsf{T}}$, where the columns of $U_b$ are eigenvectors and $d_{bi}$ is the $i$th eigenvalue. After eigenvalue flooring and ridge regularization, the inverse square root for component $b$ was

$$W_b = U_b \operatorname{diag}\left[\{\max(d_{bi}, 10^{-8}) + 10^{-2}\}^{-1/2}\right] U_b^{\mathsf{T}}. \tag{S5}$$

Here, $10^{-8}$ is the eigenvalue floor and $10^{-2}$ is the ridge term. The component-specific matrices $W_b$ were assembled into a single sparse block-diagonal transformation matrix, which was applied to the response, the intercept column, the focal accessibility predictor, and the count-based baseline covariate. Each model was then fitted by ordinary least squares using the transformed variables, without adding an additional intercept. Model-based coefficient standard errors were obtained from the transformed fit. Nominal one-sided $P$-values for $\beta_k > 0$ were calculated from the upper tail of a $t$ reference distribution using the coefficient's $t$ statistic and the residual degrees of freedom.

#### S1.7 Peak-level association models

For peak $p$, let $\mathbf{Z}_p$ denote the vector of GWAS $Z$-scores for the $n_p$ retained SNPs within that peak, and let $R_p$ denote their reference-panel LD matrix. We first symmetrized $R_p$ and computed its eigendecomposition $R_p = U_p \operatorname{diag}(d_p) U_p^{\mathsf{T}}$, where the columns of $U_p$ are eigenvectors and $d_{pi}$ is the $i$th eigenvalue. We defined the ridge-regularized inverse as

$$\Omega_p = U_p \operatorname{diag}\left[\{\max(d_{pi}, 10^{-8}) + 10^{-2}\}^{-1}\right] U_p^{\mathsf{T}} \tag{S6}$$

and summarized within-peak GWAS association by

$$Y_p = \frac{1}{n_p} \mathbf{Z}_p^{\mathsf{T}} \Omega_p \mathbf{Z}_p. \tag{S7}$$

Let $C_{pc}$ denote the ATAC count for peak $p$ in cell $c$, and let $s_c$ denote the library-size factor defined in the SNP-level analysis. For cell type $k$, the peak-level count-based accessibility and the baseline accessibility were defined as

$$A_{kp} = \frac{1}{N_k} \sum_{c \in \mathcal{C}_k} \frac{C_{pc}}{s_c}, \qquad A_p = \frac{1}{N} \sum_{c=1}^{N} \frac{C_{pc}}{s_c}. \tag{S8}$$

The winsorized peak-level WRS score $Z_{kp}^{\mathrm{WRS}}$ was obtained as described in Section S1.5. The same count-based baseline covariate $A_p$ was retained in both peak-level specifications. For each cell-type group $k$, the peak-level models were

$$Y_p = \beta_0 + \beta_k A_{kp} + \gamma_k A_p + \epsilon_p, \tag{S9}$$

$$Y_p = \beta_0 + \beta_k Z_{kp}^{\mathrm{WRS}} + \gamma_k A_p + \epsilon_p. \tag{S10}$$

Coefficients were estimated separately for each specification. The term $\epsilon_p$ denotes residual variation in $Y_p$ not explained by the focal accessibility predictor and the count-based baseline covariate.

For peaks $p$ and $q$, let $R_{pq}$ denote the reference-panel genotype-correlation matrix between their SNP sets. Under a mean-zero multivariate normal null model, the covariance of the GWAS $Z$-scores was specified by the corresponding reference-panel genotype correlations. Define $\widetilde{\mathbf{Z}}_p = \Omega_p^{1/2} \mathbf{Z}_p$ and $\widetilde{\mathbf{Z}}_q = \Omega_q^{1/2} \mathbf{Z}_q$, using the symmetric square roots of the regularized inverse matrices defined above. Their joint covariance has the block form

$$\operatorname{Cov}\left[\begin{pmatrix} \widetilde{\mathbf{Z}}_p \\ \widetilde{\mathbf{Z}}_q \end{pmatrix}\right] = \begin{pmatrix} \Omega_p^{1/2} R_p \Omega_p^{1/2} & M_{pq} \\ M_{pq}^{\mathsf{T}} & \Omega_q^{1/2} R_q \Omega_q^{1/2} \end{pmatrix}, \tag{S11}$$

where $M_{pq} = \Omega_p^{1/2} R_{pq} \Omega_q^{1/2}$. Because the inverse matrices include regularization, the diagonal blocks are not generally identity matrices. Since $Y_p = n_p^{-1} \widetilde{\mathbf{Z}}_p^{\mathsf{T}} \widetilde{\mathbf{Z}}_p$, the working covariance for peak pairs on the same chromosome with no more than 5 Mb between their genomic intervals was calculated as

$$\Sigma_{pq}^{(\mathrm{peak})} = \operatorname{Cov}(Y_p, Y_q) = \frac{2 \operatorname{tr}(M_{pq} M_{pq}^{\mathsf{T}})}{n_p n_q}. \tag{S12}$$

The diagonal entries were obtained by setting $q = p$, giving $\Sigma_{pp}^{(\text{peak})} = \text{Var}(Y_p) = 2\,\text{tr}(M_{pp}^2)/n_p^2$. Off-diagonal entries for peaks on different chromosomes or separated by more than 5 Mb were set to zero. Let $\mathbf{\Sigma}^{(\text{peak})} = (\Sigma_{pq}^{(\text{peak})})_{p,q=1}^m$ denote the resulting working covariance matrix for the $m$ retained peaks. Nonzero off-diagonal entries defined edges between peaks, yielding $B$ connected components, with isolated peaks treated as singleton components. With peaks ordered by component, the matrix can be written as

$$\mathbf{\Sigma}^{(\text{peak})} = \begin{pmatrix} \mathbf{\Sigma}_1^{(\text{peak})} & \mathbf{0} & \cdots & \mathbf{0} \\ \mathbf{0} & \mathbf{\Sigma}_2^{(\text{peak})} & \cdots & \mathbf{0} \\ \vdots & \vdots & \ddots & \vdots \\ \mathbf{0} & \mathbf{0} & \cdots & \mathbf{\Sigma}_B^{(\text{peak})} \end{pmatrix}, \tag{S13}$$

where $\mathbf{\Sigma}_b^{(\text{peak})}$ is the covariance submatrix for the peaks in component $b$. For each component, a ridge-stabilized inverse square root was computed using the same eigenvalue floor and ridge term as in the SNP-level analysis. These component-specific matrices were assembled into a single sparse transformation matrix, which was applied to the response, the intercept column, the focal accessibility predictor, and the count-based baseline covariate before ordinary least-squares fitting. Both specifications tested $H_0 : \beta_k \leq 0$ against $H_1 : \beta_k > 0$.

For each of the four component models, nominal one-sided $P$-values for the focal coefficient were adjusted using the Benjamini–Hochberg (BH) procedure separately within each trait across the tested cell-type groups. The Cauchy combination test used the nominal one-sided $P$-values rather than the multiplicity-adjusted values.

##### S1.8 Cross-resolution evidence combination

For each combination of a trait and a cell type, all available nominal one-sided $P$-values from the SNP-count, SNP-WRS, peak-count, and peak-WRS models were combined with equal weights using the Cauchy combination test (Liu and Xie, 2020; Gui et al., 2025).

The resulting nominal omnibus $P$-values were adjusted using the Benjamini–Hochberg procedure within each GWAS dataset across the tested cell-type groups. Component-level results were retained to aid interpretation of the omnibus results. Statistical significance was defined as a within-trait BH-adjusted $P < 0.05$.

##### S1.9 Complementary S-LDSC analysis

We used S-LDSC (Finucane et al., 2015) as a complementary annotation-level analysis for cell-type prioritization. For each cell-type group, the detection fraction for each post-QC autosomal peak was defined as the proportion of cells with nonzero peak counts. Detection fold changes relative to all other cells were calculated as the ratio of the within-group and remaining-cell detection fractions after adding $10^{-4}$ to each. Peaks were selected if the within-group detection fraction was at least 1% and the detection fold change was at least 2. Peaks were ranked in descending order of the product of $\log_2$ detection fold change and within-group detection fraction. If more than 50,000 peaks passed the thresholds, the 50,000 highest-ranked peaks were retained. If fewer than 500 peaks passed, the top 5000 ranked peaks from all retained peaks were used instead. Selected peak intervals were merged to construct binary SNP annotations. Annotation-specific LD scores were calculated using a 1-cM window.

Each custom annotation was tested separately in an S-LDSC model that included baselineLD v2.2 (Gazal et al., 2017). The analysis used LD-unpruned GWAS summary statistics, LD scores based on 1000 Genomes Phase 3 European reference genotypes, and HapMap3 non-MHC regression SNPs (International HapMap 3 Consortium, 2010). Annotation coefficient estimates and their $z$-scores were extracted, together with heritability-enrichment estimates and their $P$-values. For cell-type prioritization, one-sided $P$-values for positive annotation coefficients were calculated from the coefficient $z$-scores using the upper tail of the standard normal distribution. These $P$-values were adjusted across 24 cell-type groups within each GWAS dataset using the Benjamini–Hochberg procedure. S-LDSC was interpreted as a complementary annotation-level analysis rather than independent validation of the SNP- and peak-level models.

##### S1.10 Implementation

The primary workflow was implemented in R with Seurat, Signac, GenomicRanges, SNPRelate, presto, Matrix, igraph, and BiocParallel; S-LDSC was run separately with its Python command-line implementation. The implementation uses sparse matrix representations and supports parallel SNP-accessibility quantification across feature chunks through BiocParallel. Intermediate objects and model results were saved as RDS files after GWAS processing, accessibility quantification, LD covariance construction, and model fitting, allowing individual stages to be rerun using saved upstream outputs. A fixed random seed was used for LD pruning. The EPIC2 source code, documentation, and an example workflow with test data are available at `https://github.com/LingxuanZ/EPIC2`. Scripts for reproducing the analyses reported in this study are available at `https://github.com/LingxuanZ/EPIC2/tree/main/manuscript_reproducibility`.

#### S2 Supplementary Results

##### S2.1 Fetal atlas and trait-specific accessibility inputs

The starting scATAC-seq object contained 87,526 cells from seven fetal tissues and a shared set of 1,050,819 ATAC peaks. Cell-level filtering retained 84,238 cells, and peak-level filtering retained 915,345 peaks, corresponding to 96.2% and 87.1% of the starting cells and peaks, respectively. The retained cells were visualized using UMAP and colored by tissue and cell-type annotations (Supplementary Fig. S1).

After LD pruning and removal of strand-ambiguous SNPs, the processed GWAS inputs contained 348,323–634,030 SNPs for the psychiatric GWAS datasets and 395,305–396,599 SNPs for the lipid GWAS datasets. Of these, 56,262–99,447 SNPs per psychiatric GWAS dataset and 64,917–65,156 SNPs per lipid GWAS dataset overlapped the filtered fetal ATAC peaks. After removing SNPs with zero total fragment coverage across all cells, the final SNP-by-cell matrices contained 56,262–99,445 SNPs for the psychiatric GWAS datasets and 64,917–65,156 SNPs for the lipid GWAS datasets (Supplementary Table S1). Corresponding peak-by-cell matrices were constructed from filtered ATAC peaks overlapping at least one retained SNP.

##### S2.2 LD-aware associations and complementary analyses

As expected from one-versus-rest WRS feature selection, the top-ranked SNP and peak features generally showed preferential accessibility in the focal group, although some were also accessible in other groups (Supplementary Figs. S2 and S3).

Supplementary Table S2 reports nominal $P$-values from the four component tests and the Cauchy combination test for the 12 associations identified by EPIC2 after within-trait BH correction ($P_{\text{adj}} < 0.05$). All 12 associations were also significant in the peak-WRS model after within-trait

BH correction. For TC in liver hepatoblasts, the peak-WRS model yielded the smallest nominal and BH-adjusted $P$-values among the four component tests ($P = 1.43 \times 10^{-5}$, $P_{\text{adj}} = 2.25 \times 10^{-4}$). For nine of the 12 associations, a peak-level component yielded the smallest nominal $P$-value among the four component tests. These results highlight the contribution of peak-level representations and support examining component results alongside the omnibus test.

EPIC2's Cauchy combination test identified 12 significant associations between traits and cell types across six GWAS datasets after within-trait BH correction: three each for LDL and TC, two each for TG and SCZBIP, and one each for SCZ and SCZ2. No associations for HDL or BIP were significant after BH correction. Lipid associations involved liver hepatoblasts (LDL, TC, and TG), liver myeloid cells (LDL and TC), kidney ureteric bud cells (LDL and TC), and kidney vascular endothelial cells (TG). The four psychiatric associations involved cerebrum excitatory neurons for SCZ, SCZ2, and SCZBIP, and cerebrum inhibitory neurons for SCZBIP, with the strongest omnibus evidence observed for SCZ in excitatory neurons ($P = 5.40 \times 10^{-6}$, $P_{\text{adj}} = 1.30 \times 10^{-4}$). In the S-LDSC analysis, three associations between traits and cell types were significant after within-trait BH correction using one-sided tests for positive annotation coefficients. Both EPIC2 and S-LDSC prioritized cerebrum excitatory neurons for SCZ and SCZ2, whereas adrenal chromaffin cells were prioritized for SCZ2 only by S-LDSC. The S-LDSC coefficient for SCZBIP in cerebrum excitatory neurons was nominally significant ($P = 0.00272$), but was not significant after BH correction ($P_{\text{adj}} = 0.0654$). All associations between lipid traits and cell types identified by EPIC2 had positive S-LDSC annotation coefficients, but none were significant after BH correction in S-LDSC (Supplementary Table S3).

### S3 Supplementary Discussion

This study presents an LD-aware, cross-resolution framework for relating GWAS association magnitude to quantitative chromatin accessibility. GREGOR, GARFIELD, and RLEA evaluate enrichment using variant sets or GWAS significance thresholds and overlap with regulatory annotations (Schmidt et al., 2015; Iotchkova et al., 2019; Rosean et al., 2024). In contrast, the primary EPIC2 models retain continuous GWAS association statistics and quantitative accessibility among the analyzed variants. At each resolution, EPIC2 evaluates count-based accessibility and one-versus-rest WRS scores in separate models. In their published workflows, g-chromVAR and SCAVENGE use fine-mapped variant posterior probabilities to weight chromatin features, and SCAVENGE additionally propagates trait-relevance information over a cell–cell graph (Ulirsch et al., 2019; Yu et al., 2022). The EPIC2 association tests do not require fine-mapping posterior probabilities or cell–cell graph propagation. The SNP-level models retain information at variant positions, whereas the peak-level models summarize association strength and accessibility across regulatory elements; both incorporate local LD-based working covariance matrices. The workflow combines the component $P$-values using the Cauchy test and retains the component statistics to help identify which representations support an association.

The lipid results showed partial agreement with established tissue biology. EPIC2 prioritized liver hepatoblasts for LDL, TC, and TG, and the TC association was the only Cauchy hit supported by all four component tests after within-trait BH correction. An integrative analysis of lipid GWAS and adult human and mouse liver scRNA-seq data reported enrichment of lipid GWAS signals in gene sets associated with periportal hepatocytes (Hao et al., 2020), while a human liver developmental atlas reconstructed a developmental trajectory from hepatoblasts through fetal hepatocytes to adult hepatocytes (Wesley et al., 2022). The liver findings are therefore coherent at the tissue and lineage levels, although fetal hepatoblasts should not be equated with mature hepatocytes. The absence of an HDL Cauchy hit does not exclude hepatic involvement and may reflect limited statistical power, developmental specificity, or the restricted tissue coverage of this atlas.

Additional lipid associations involved kidney ureteric bud cells and vascular endothelial cells. The collecting duct lineage provides a possible context for the ureteric bud associations with LDL and TC. Human ureteric bud organoids can differentiate into functional collecting duct cells (Shi et al., 2023). Collecting duct-derived PCSK9 has been implicated in hypercholesterolemia in nephrotic syndrome. In mice treated with nephrotoxic serum, collecting duct-specific *Pcsk9* deficiency prevented hypercholesterolemia (Molina-Jijon et al., 2020). In nephrotic syndrome models, circulating ANGPTL4 reduced proteinuria by binding to glomerular endothelial $\alpha_v\beta_5$ integrin. It also promoted hypertriglyceridemia by inhibiting lipoprotein lipase in skeletal muscle, heart, and adipose tissue (Clement et al., 2014). These findings provide a disease-related context for considering the kidney vascular endothelial association with TG. However, they do not establish that kidney endothelial cells mediate TG variation. The PCSK9 and ANGPTL4 studies examined nephrotic disease, not the fetal cell populations analyzed here. None of the lipid associations prioritized by EPIC2 were BH-significant in S-LDSC, although all corresponding annotation coefficients were positive.

The psychiatric findings were consistent with prior cellular evidence. EPIC2 prioritized cerebrum excitatory neurons for SCZ, SCZ2, and SCZBIP, and the SCZ and SCZ2 associations were also BH-significant in S-LDSC. This pattern is consistent with SCZ heritability enrichment near genes specifically expressed in excitatory neurons (Finucane et al., 2018), enrichment of SCZ risk variants in glutamatergic-neuron open chromatin (Hauberg et al., 2020), and enrichment of SCZ genetic risk in genes preferentially expressed in developing prenatal neurons (Cameron et al., 2023). EPIC2 additionally prioritized cerebrum inhibitory neurons for SCZBIP, although this association was not significant after BH correction in S-LDSC. The SCZBIP findings are also compatible with the neuronal and synaptic pathways reported in the source GWAS (Ruderfer et al., 2018), although the combined case–control analysis does not establish that either neuronal association is shared by both disorders.

The psychiatric datasets share contributing cohorts, so recurrence of the excitatory-neuron signal represents cross-dataset consistency rather than independent replication. Similarly, the absence of a BH-significant BIP association should not be interpreted as evidence against neuronal involvement. Previous work reported BIP heritability enrichment near genes specifically expressed in GABAergic neurons (Finucane et al., 2018), while a larger BIP GWAS implicated genes with high expression specificity in neurons of the prefrontal cortex and hippocampus (Mullins et al., 2021). Statistical power, developmental stage, and cell-type classification may therefore influence which cell types are prioritized.

Several limitations constrain interpretation. First, EPIC2 prioritizes trait-relevant cellular contexts but does not establish causal regulatory mechanisms. It does not determine which variants are causal, how individual alleles affect chromatin accessibility, or which genes mediate the associations. Peak-level aggregation can combine association evidence from multiple variants within the same peak without identifying their individual regulatory roles. Second, the present application was restricted to seven fetal tissues and did not assess donor-level replication, limiting generalization across developmental stages, individuals, and unobserved cell states. The restriction to fetal data is especially relevant to adult lipid phenotypes. Third, results may depend on LD pruning, the local covariance window, ridge regularization, and the suitability of the reference panel. Null calibration and robustness to covariance misspecification warrant further evaluation under realistic LD and accessibility structures. Sparse accessibility, peak calling, group size, and correlated accessibility profiles may also affect component results and limit attribution of a signal to a single cell type. Fourth, a Cauchy result can be driven by one strong component, making the retained component statistics important for interpretation; the combination does not compensate for miscalibrated component $P$-values (Gui et al., 2025). S-LDSC provides an annotation-level benchmark for EPIC2 in this application; agreement between the methods reflects concordance across analyses of shared data rather than independent replication.

Overall, EPIC2 provides a framework for cell-type prioritization from GWAS summary statistics and chromatin accessibility data without requiring fine-mapping posterior probabilities. The observed liver and excitatory-neuron associations support the biological plausibility of the prioritizations in this application, while additional method-specific findings provide candidates for follow-up. Evaluation in broader multi-donor fetal and adult atlases and ancestry-matched settings would help assess generalizability. Integration with fine-mapping, allele-specific and peak-to-gene evidence, and functional experiments could help connect prioritized cellular contexts to regulatory mechanisms.

### S4 Supplementary Figures

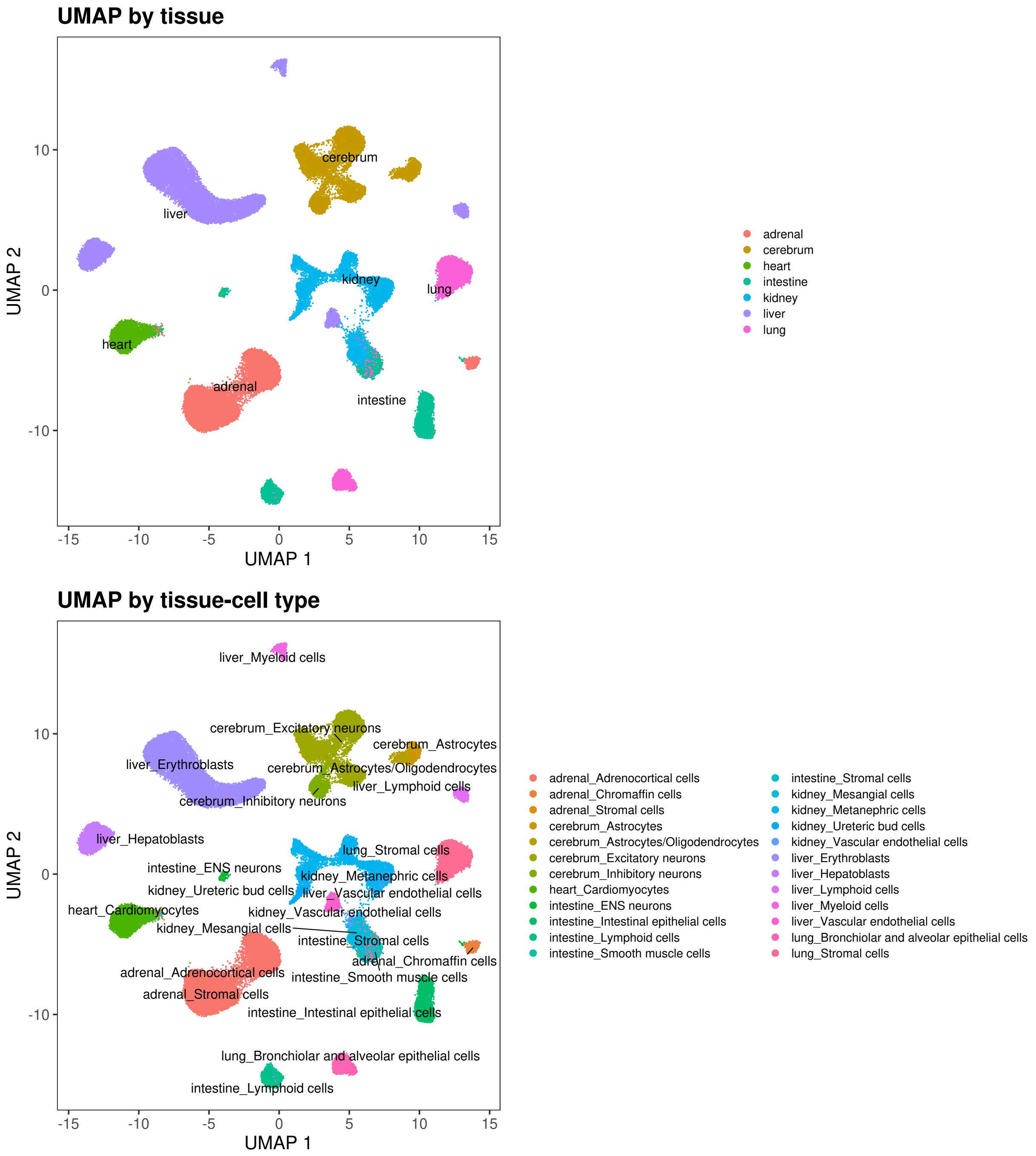

Figure S1: UMAP visualization of the analyzed subset of the human fetal single-cell ATAC-seq atlas after cell- and peak-level quality control. Each point represents one retained cell. Both panels show the same UMAP embedding, with cells colored by tissue (top) or cell-type group (bottom), based on annotations from the atlas.

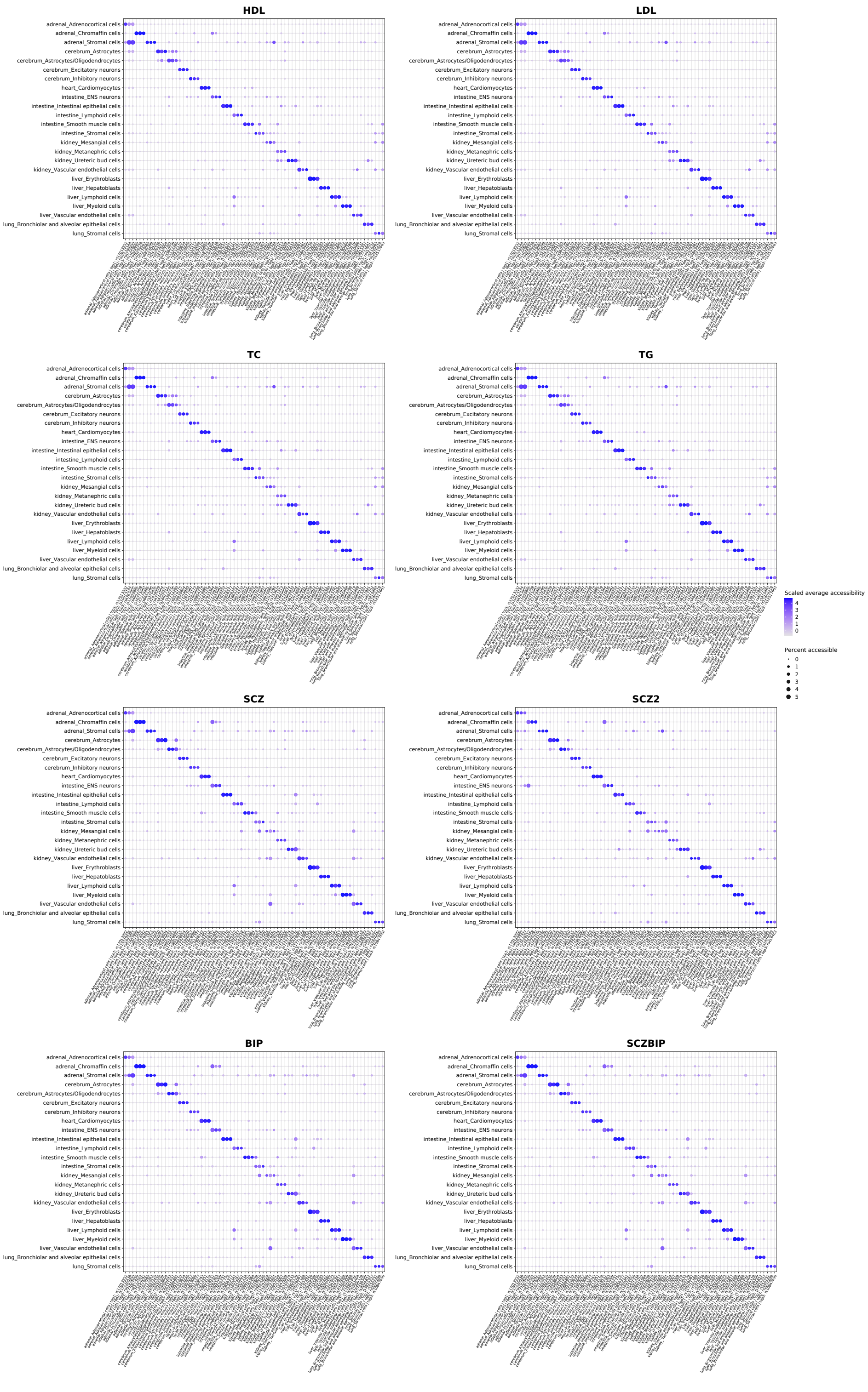

Figure S2: Accessibility patterns of the top-ranked SNPs from one-sided Wilcoxon rank-sum (WRS) tests across fetal cell-type groups. For each combination of a trait and a cell type, up to three retained SNP positions with AUC $\geq 0.5$ and the smallest nominal one-sided WRS $P$-values were selected, and their accessibility was visualized across all groups. Dot size represents the percentage of cells in each group with nonzero accessibility at the SNP. Color represents the group mean accessibility, transformed as $\log(1 + x)$ and standardized across groups within each SNP. WRS $P$-values were used for SNP selection and are not encoded by dot size or color.

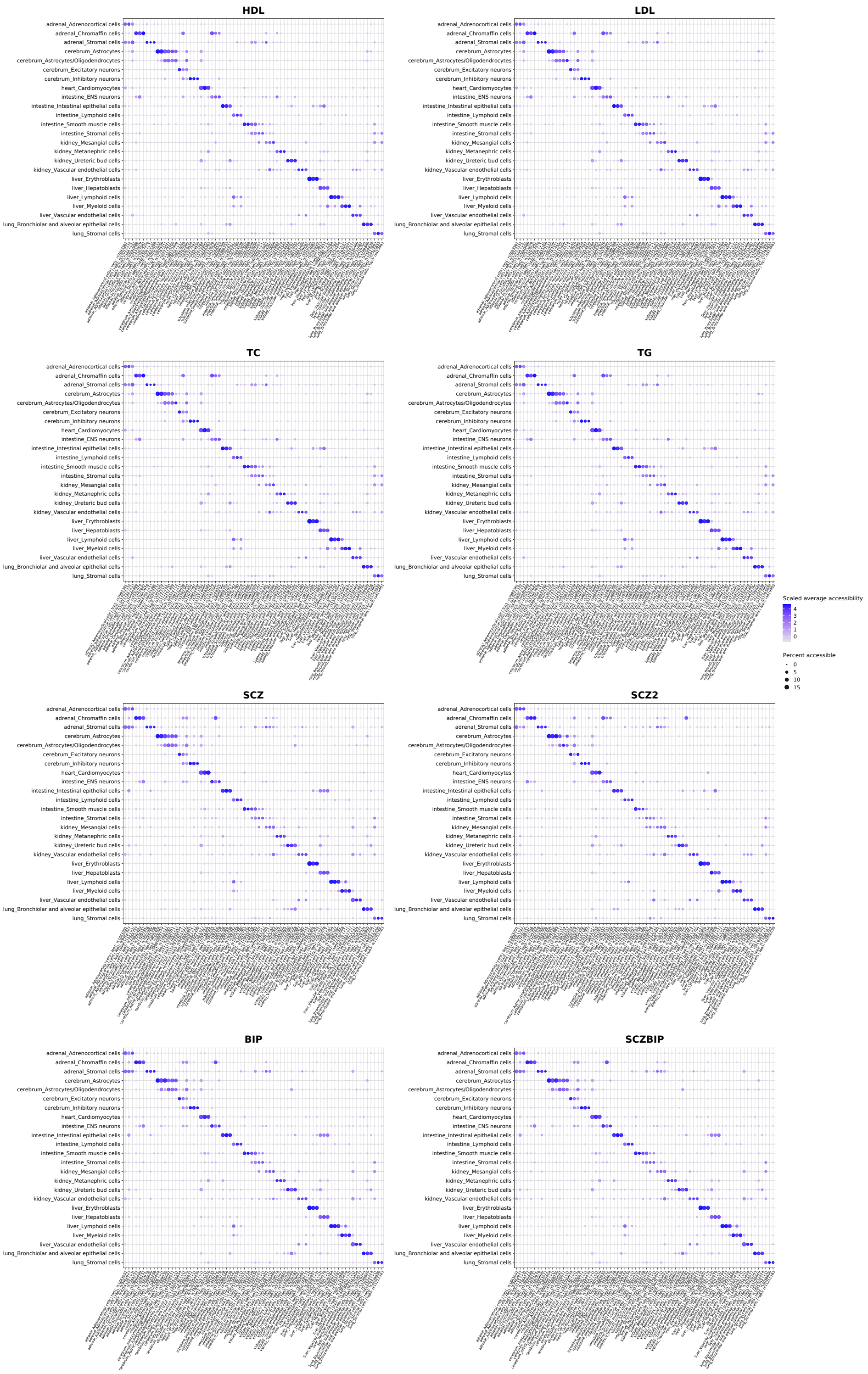

Figure S3: Accessibility patterns of the top-ranked GWAS-overlapping ATAC peaks from one-sided Wilcoxon rank-sum (WRS) tests across fetal cell-type groups. For each combination of a trait and a cell type, up to three peaks with AUC $\geq 0.5$ and the smallest nominal one-sided WRS $P$-values were selected, and their accessibility was visualized across all groups. Dot size represents the percentage of cells in each group with nonzero accessibility at the peak. Color represents the group mean accessibility, transformed as $\log(1 + x)$ and standardized across groups within each peak. WRS $P$-values were used for peak selection and are not encoded by dot size or color.

#### S5 Supplementary Tables

Table S1: Trait-specific GWAS sources, sample sizes, and processing summary. Filtered GWAS SNPs were obtained after LD pruning and removal of strand-ambiguous variants. Final SNPs were required to have nonzero fragment coverage.

| Trait | GWAS sample size | Filtered GWAS SNPs | SNPs overlapping ATAC peaks | Final nonzero-coverage SNPs | Source |
|---|---|---|---|---|---|
| SCZ | 79,845[a] | 634,030 | 99,447 | 99,445 | Schizophrenia Working Group of the Psychiatric Genomics Consortium (2014) |
| SCZ2 | 105,318 | 348,323 | 56,262 | 56,262 | Pardiñas et al. (2018) |
| BIP | 51,710 | 630,665 | 99,033 | 99,031 | Stahl et al. (2019) |
| SCZBIP | 107,620 | 630,725 | 99,016 | 99,014 | Ruderfer et al. (2018) |
| HDL | 188,577[b] | 396,572 | 65,150 | 65,150 | Willer et al. (2013) |
| LDL | 188,577[b] | 395,305 | 64,917 | 64,917 | Willer et al. (2013) |
| TC | 188,577[b] | 396,599 | 65,156 | 65,156 | Willer et al. (2013) |
| TG | 188,577[b] | 395,414 | 64,940 | 64,940 | Willer et al. (2013) |

[a]Case–control participants only; the primary PGC GWAS also included 1,235 trios comprising an affected offspring and both parents.

[b]Study-level sample size reported in the published article; variant-specific sample sizes are provided in the released summary statistics and vary across variants.

Table S2: LD-aware association results across fetal cell-type groups. Results from the four one-sided component tests and the Cauchy omnibus test for the 12 associations between traits and cell types with within-trait BH-adjusted Cauchy $P < 0.05$. All $P$-values shown are nominal, including the Cauchy combined $P$-values in the final column. The Cauchy test combines the four nominal component $P$-values with equal weights. Feature counts indicate the numbers of final nonzero-coverage SNPs and peak-level observations, respectively. BH, Benjamini–Hochberg; WRS, Wilcoxon rank-sum.

| Trait | Cell type | Features SNP/peak | SNP count $P$ | SNP WRS $P$ | Peak count $P$ | Peak WRS $P$ | Cauchy-combined $P$ |
|---|---|---|---|---|---|---|---|
| LDL | liver: Hepatoblasts | 64,917 / 59,337 | $3.00 \times 10^{-2}$ | $3.69 \times 10^{-2}$ | $8.77 \times 10^{-5}$ | $1.81 \times 10^{-4}$ | $2.36 \times 10^{-4}$ |
| LDL | kidney: Ureteric bud cells | 64,917 / 59,337 | $4.62 \times 10^{-2}$ | $5.45 \times 10^{-2}$ | $5.03 \times 10^{-4}$ | $1.69 \times 10^{-3}$ | $1.53 \times 10^{-3}$ |
| LDL | liver: Myeloid cells | 64,917 / 59,337 | $1.58 \times 10^{-3}$ | $4.84 \times 10^{-2}$ | $3.12 \times 10^{-2}$ | $4.89 \times 10^{-3}$ | $4.49 \times 10^{-3}$ |
| TC | liver: Hepatoblasts | 65,156 / 59,520 | $3.44 \times 10^{-4}$ | $8.86 \times 10^{-4}$ | $4.27 \times 10^{-5}$ | $1.43 \times 10^{-5}$ | $4.10 \times 10^{-5}$ |
| TC | kidney: Ureteric bud cells | 65,156 / 59,520 | $4.80 \times 10^{-2}$ | $4.11 \times 10^{-2}$ | $1.42 \times 10^{-4}$ | $1.88 \times 10^{-5}$ | $6.63 \times 10^{-5}$ |
| TC | liver: Myeloid cells | 65,156 / 59,520 | $4.12 \times 10^{-3}$ | $2.06 \times 10^{-2}$ | $1.44 \times 10^{-2}$ | $1.31 \times 10^{-3}$ | $3.56 \times 10^{-3}$ |
| TG | kidney: Vascular endothelial cells | 64,940 / 59,358 | $1.10 \times 10^{-2}$ | $1.13 \times 10^{-4}$ | $1.33 \times 10^{-2}$ | $6.31 \times 10^{-4}$ | $3.76 \times 10^{-4}$ |
| TG | liver: Hepatoblasts | 64,940 / 59,358 | $2.60 \times 10^{-2}$ | $2.06 \times 10^{-4}$ | $5.62 \times 10^{-2}$ | $2.70 \times 10^{-4}$ | $4.64 \times 10^{-4}$ |
| SCZ | cerebrum: Excitatory neurons | 99,445 / 84,869 | $1.35 \times 10^{-1}$ | $3.83 \times 10^{-4}$ | $1.45 \times 10^{-1}$ | $1.35 \times 10^{-6}$ | $5.40 \times 10^{-6}$ |
| SCZ2 | cerebrum: Excitatory neurons | 56,262 / 50,898 | $2.79 \times 10^{-2}$ | $3.16 \times 10^{-3}$ | $1.16 \times 10^{-1}$ | $5.00 \times 10^{-4}$ | $1.69 \times 10^{-3}$ |
| SCZBIP | cerebrum: Excitatory neurons | 99,014 / 84,503 | $7.47 \times 10^{-2}$ | $2.81 \times 10^{-3}$ | $1.40 \times 10^{-1}$ | $1.45 \times 10^{-5}$ | $5.77 \times 10^{-5}$ |
| SCZBIP | cerebrum: Inhibitory neurons | 99,014 / 84,503 | $7.03 \times 10^{-3}$ | $7.71 \times 10^{-3}$ | $5.67 \times 10^{-2}$ | $7.57 \times 10^{-4}$ | $2.48 \times 10^{-3}$ |

Table S3: Comparison of associations between traits and cell types with BH-adjusted $P < 0.05$ in either the EPIC2 Cauchy omnibus analysis or the S-LDSC annotation-coefficient analysis. Nominal and BH-adjusted $P$-values are shown for both methods, together with the S-LDSC annotation-coefficient $z$-statistic. S-LDSC $P$-values are from one-sided tests for positive annotation coefficients. BH correction was applied separately for each method across 24 cell-type groups within each GWAS dataset. Classification indicates whether an association was significant in EPIC2, S-LDSC, or both.

| Trait | Cell type | EPIC2 Cauchy-combined $P$ | EPIC2 BH-adjusted $P$ | S-LDSC coefficient $z$ | S-LDSC $P$ | S-LDSC BH-adjusted $P$ | Classification |
|---|---|---|---|---|---|---|---|
| LDL | liver: Hepatoblasts | $2.36 \times 10^{-4}$ | $5.65 \times 10^{-3}$ | $1.84 \times 10^{0}$ | $3.31 \times 10^{-2}$ | $4.02 \times 10^{-1}$ | EPIC2 only |
| LDL | kidney: Ureteric bud cells | $1.53 \times 10^{-3}$ | $1.83 \times 10^{-2}$ | $5.19 \times 10^{-1}$ | $3.02 \times 10^{-1}$ | $6.58 \times 10^{-1}$ | EPIC2 only |
| LDL | liver: Myeloid cells | $4.49 \times 10^{-3}$ | $3.59 \times 10^{-2}$ | $1.04 \times 10^{0}$ | $1.50 \times 10^{-1}$ | $5.13 \times 10^{-1}$ | EPIC2 only |
| TC | liver: Hepatoblasts | $4.10 \times 10^{-5}$ | $7.96 \times 10^{-4}$ | $2.46 \times 10^{0}$ | $6.89 \times 10^{-3}$ | $1.65 \times 10^{-1}$ | EPIC2 only |
| TC | kidney: Ureteric bud cells | $6.63 \times 10^{-5}$ | $7.96 \times 10^{-4}$ | $9.20 \times 10^{-1}$ | $1.79 \times 10^{-1}$ | $4.29 \times 10^{-1}$ | EPIC2 only |
| TC | liver: Myeloid cells | $3.56 \times 10^{-3}$ | $2.85 \times 10^{-2}$ | $1.21 \times 10^{0}$ | $1.12 \times 10^{-1}$ | $3.85 \times 10^{-1}$ | EPIC2 only |
| TG | kidney: Vascular endothelial cells | $3.76 \times 10^{-4}$ | $5.57 \times 10^{-3}$ | $6.87 \times 10^{-1}$ | $2.46 \times 10^{-1}$ | $9.09 \times 10^{-1}$ | EPIC2 only |
| TG | liver: Hepatoblasts | $4.64 \times 10^{-4}$ | $5.57 \times 10^{-3}$ | $6.27 \times 10^{-1}$ | $2.65 \times 10^{-1}$ | $9.09 \times 10^{-1}$ | EPIC2 only |
| SCZ | cerebrum: Excitatory neurons | $5.40 \times 10^{-6}$ | $1.30 \times 10^{-4}$ | $3.12 \times 10^{0}$ | $9.01 \times 10^{-4}$ | $2.16 \times 10^{-2}$ | Both EPIC2 and S-LDSC |
| SCZ2 | cerebrum: Excitatory neurons | $1.69 \times 10^{-3}$ | $4.07 \times 10^{-2}$ | $3.28 \times 10^{0}$ | $5.16 \times 10^{-4}$ | $1.24 \times 10^{-2}$ | Both EPIC2 and S-LDSC |
| SCZ2 | adrenal: Chromaffin cells | $8.17 \times 10^{-2}$ | $2.80 \times 10^{-1}$ | $2.91 \times 10^{0}$ | $1.79 \times 10^{-3}$ | $2.15 \times 10^{-2}$ | S-LDSC only |
| SCZBIP | cerebrum: Excitatory neurons | $5.77 \times 10^{-5}$ | $1.38 \times 10^{-3}$ | $2.78 \times 10^{0}$ | $2.72 \times 10^{-3}$ | $6.54 \times 10^{-2}$ | EPIC2 only |
| SCZBIP | cerebrum: Inhibitory neurons | $2.48 \times 10^{-3}$ | $2.98 \times 10^{-2}$ | $1.81 \times 10^{0}$ | $3.55 \times 10^{-2}$ | $3.14 \times 10^{-1}$ | EPIC2 only |
